# Adaptation across a precipitation gradient from niche center to niche edge

**DOI:** 10.1101/2024.04.15.589238

**Authors:** Samantha vanDeurs, Oliver Reutimann, Hirzi Luqman, Dikla Lifshitz, Einav Mayzlish Gati, Jake Alexander, Simone Fior

## Abstract

Evaluating the potential for species to adapt to changing climate relies on an appreciation of current patterns of adaptive variation and selection, which might vary in intensity across a species’ niche, affecting our inference of where adaptation might be most important in the future. Here we investigate the genetic basis of adaptation in *Lactuca serriola*, the wild relative of the common lettuce, growing along a steep precipitation gradient in Israel approaching the species’ arid niche limit, and used candidate loci to inform predictions of its past and future adaptive evolution. Environmental association analyses combined with generalized dissimilarity models revealed 98 candidate genes showing non-linear shifts in allele frequencies across the gradient, with a large proportion under strong directional selection near the dry niche edge. Selection acts on genes with separate suites of biological functions, specifically related to osmotic stress and phenological adjustments at the dry edge, and related to biotic interactions at the niche center. The adaptive genetic composition of populations, as inferred through polygenic risk scores, indicates that intensified selection may be the key determinant of the dry niche edge. However, inference of past and future evolutionary change suggests larger adaptive shifts in the mesic part of the range, which is most affected by climate change. Our study highlights that non-linear spatial variation in selection has implications for predicting past and future responses to climate change.

## Introduction

While many species are migrating to track their historical climate, many more are failing to match the current pace of climate change and so are being exposed to novel environmental conditions (Alexander et al., 2018; Corlett & Westcott, 2013). Adaptive evolution may help some species to persist *in situ* under changing climates (Anderson et al., 2012), as evidenced by the existence of local climate adaptation in populations today (Halbritter et al., 2018; Leimu & Fischer, 2008). However, the need for adaptation to maintain population persistence is expected to vary across a species’ range, due to spatial variation in both the magnitude of climatic changes, and/or the intensity of selection (Exposito-Alonso et al., 2019). In particular, selection intensity is expected to vary from weaker stabilizing selection under more benign climatic conditions near the niche center, to stronger directional selection under more extreme environmental conditions towards the niche edge (Angert et al., 2020; Brown, 1984; Hargreaves et al., 2014; Lee-Yaw et al., 2016). Furthermore, climatic changes are not expected to be uniform, potentially resulting in heterogeneous shifts in selective pressures (Ban et al., 2015). Closer to the niche edge, populations are likely at or near their physiological limits due to extreme climatic events like frost, heatwaves, or drought driving conditions far from the species’ optimum (Gaston, 2003; Hargreaves et al., 2014). These edge populations, often the first to confront new environmental stressors, might also lack pre-adapted genotypes within the species’ gene pool (Nadeau & Urban, 2019) or face other genetic constraints to adaptation (Bridle & Vines, 2007; Sexton et al., 2009). Because plasticity alone is unlikely to allow populations to persist under novel conditions in the long-term (Visser, 2008), they are potentially the first populations facing the risk of extinction under a changing climate (Wessely et al., 2022). Investigating patterns of selection across the species’ niche combined with predictions of future climatic regimes can therefore help identify populations that are most impacted by climate change, and for which adaptive evolution might be essential for long-term persistence.

Assessing variation in both natural selection and adaptation across a species’ niche is a difficult task, since it requires evaluations of multiple populations spanning a gradient from niche center to edge (Angert et al., 2020; Etterson, 2004). Advances in genomic approaches have enhanced our ability to examine climate adaptation across the species’ niche (Fitzpatrick & Keller, 2015; Razgour et al., 2019). Compared to common garden experiments, which excel at identifying phenotypic differentiation stemming from natural selection but are logistically challenging, genomic methods offer a more nuanced, molecular understanding of climate adaptation both at individual loci and at the population level. Specifically, environmental association analyses (EAA) allow for the identification of environmental drivers of selection and genes with allelic variation underlying adaptation (Rellstab et al., 2015), and point to the biological processes of the identified genes. Moreover, as adaptation to climate often involves multiple biological functions and is thus likely to be polygenic (Barghi et al., 2020; Yeaman, 2015), evaluating the cumulative contribution of adaptive alleles to the genetic composition of populations can provide insights into the selection regimes imposed along an environmental gradient. While previous studies have linked polygenic adaptation to environmental gradients (Folkertsma et al., 2024; Xuereb et al., 2018; Yadav et al., 2021), these have not been used to examine how the strength of climatic selection may vary from niche center to edge, potentially resulting in non-linear trends of adaptation (Aguirre-Liguori et al., 2021; Fitzpatrick & Keller, 2015). Finally, projecting patterns of climate adaptation inferred across the species’ niche to future conditions constitutes our best estimate of adaptive response and persistence under climate change, particularly for niche edge populations (Aguirre-Liguori et al., 2021; Chen et al., 2023; Fitzpatrick et al., 2021).

*Lactuca serriola* is a diploid, ruderal annual plant and the most widely distributed species in the genus (Lebeda et al., 2004). *L. serriola* is the wild progenitor of the cultivated lettuce, *L. sativa*, and is notable for its drought tolerance (Werk & Ehleringer, 1985, 1986). This resilience is attributed to a root system that allows high water use efficiency, which, coupled with the species’ short generation time, allows for rapid habitat colonization and seed production under arid conditions (Chadha et al., 2019; Johnson et al., 2000). *L. serriola* is primarily self-pollinated with an estimated selfing rate of 95.5% (D’Andrea et al., 2016; Mejías, 1994) and its achenes, equipped with a pappus, facilitate effective wind-dispersal, contributing to its wide distribution (Chadha & Florentine, 2021; Mejías, 1994; Weaver & Downs, 2003). Seed germination is influenced by temperature and precipitation, suggesting that climate is a major determinant of the species’ distribution (Chadha et al., 2019; Prince et al., 1985). Originating from the Caucasus region, *L. serriola* has spread across Eurasia and has been introduced globally to all continents, with the exception of Antarctica (Kuang et al., 2008; Wei et al., 2021), occupying a large breadth of climatic conditions. This spread has been associated with climate adaptation. For example, directional selection near the cool/wet edge of the *L. serriola’s* range in northern Europe has likely driven the evolution of vernalization requirements that are lacking from populations from more arid regions (Alexander, 2013; Prince, 1980; Prince et al., 1978). In its introduced ranges, *L. serriola* has shown the ability to rapidly evolve its flowering phenology to reproduce the flowering clines observed in the native range along the gradients of aridity (Alexander, 2013), showcasing its ability to rapidly adapt to novel climates. *L. serriola* therefore possesses many properties that make it an ideal model system for studying climate adaptation.

*L. serriola* reaches its arid native range edge in the Middle Eastern deserts, including Israel’s Negev and Arava Deserts (Figure S1, S2). The country possesses a steep precipitation gradient ranging from 25 mm annually in the Arava to over 1000 mm in the Golan Heights (Goldreich, 2003). Rainfall is highly seasonal, predominantly occurring in winter, with a summer drought period with little to no precipitation, which has increased in length over the last 40 years (Drori et al., 2021). Studying *L. serriola* populations along the precipitation gradient in Israel thus provides the opportunity to examine the strength and variation of precipitation as a selective agent from the arid niche edge toward the mesic niche center (Figure S2, S3), as well as the species’ potential for adaptation to current and future precipitation regimes. Using 21 populations of *L. serriola* from Israel as our study system, we address the following questions:

1. Do populations show genomic signatures of adaptation associated with precipitation and if so, what is the genetic basis?
2. How does the intensity of selection associated with precipitation vary along the precipitation gradient from the species’ niche center to the niche edge?
3. How may the adaptive alleles of populations have shifted in response to climate change in recent decades, and what are the implications for adaptive evolution in response to future climatic scenarios?

## Materials and methods

### Population sampling and genome sequencing

We conducted all analyses in R version 4.2.3 (R Core Team, 2023). We characterized climatic conditions of *L. serriola* populations using WorldClim bioclimatic variables at a 2.5-minute resolution and selected populations to span the entirety of Israel’s precipitation gradient, with three populations falling into each 100 mm precipitation band from 100 mm to 770 mm of mean annual precipitation (Fig 1A; Fick & Hijmans, 2017)). The term mesic here (and hereafter) refers to the relatively wet conditions of Northern Israel in the context of the species’ niche as a whole, which extends into much wetter conditions (Figure S2). In August and September of 2020, we collected achenes (hereafter “seeds”) from a minimum of 15 mother plants from each of the 21 locations (hereafter “populations”).

**Figure 1:**
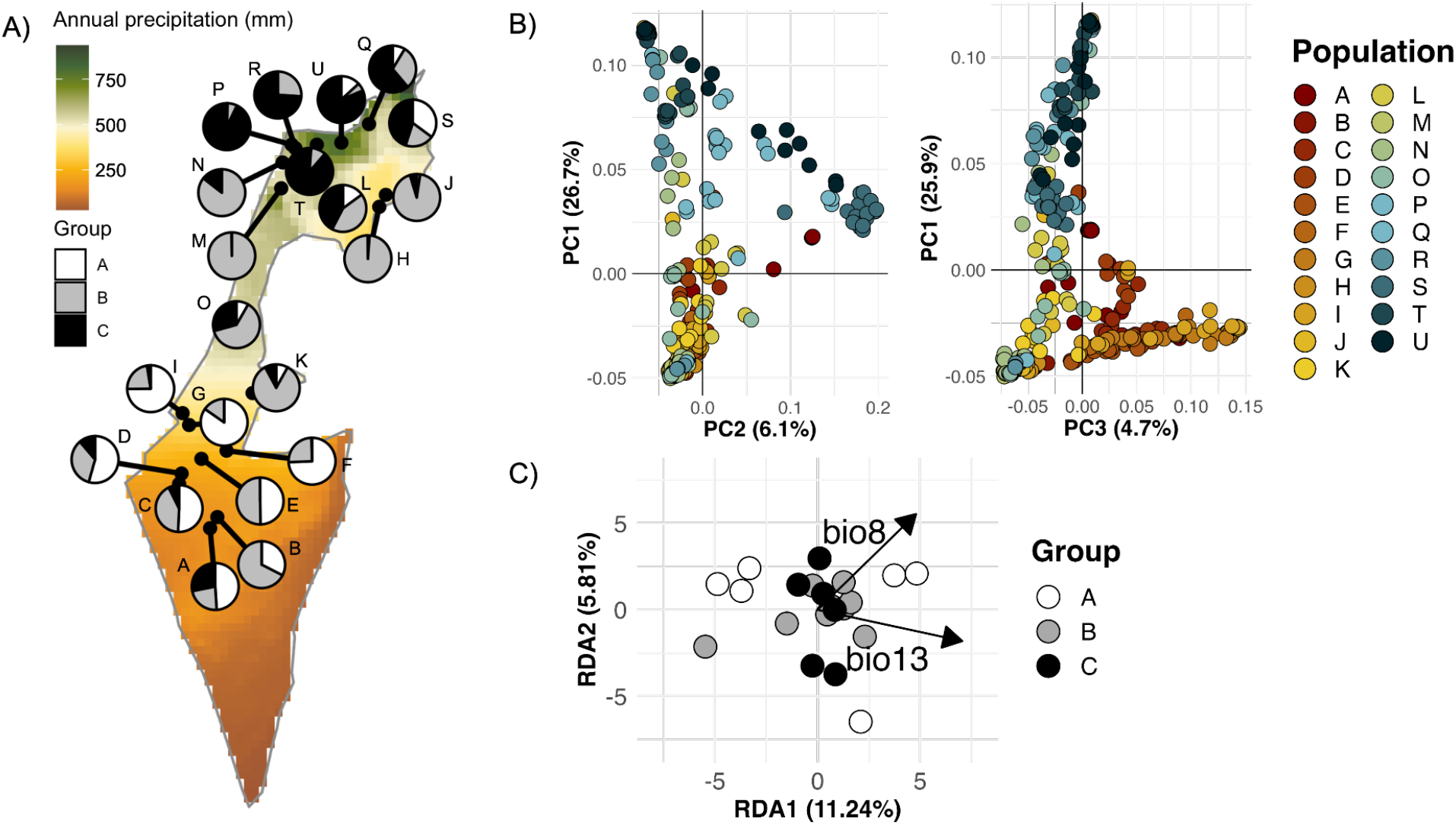
Genetic variation along the precipitation gradient in Israel. (A) Map of 21 sampled populations of *L. serriola*. Pie charts depict the average likelihood for individuals of each population to belong to one of three genomic clusters. (B) PCA of exonic genetic variation. (C) Biplot of the partial RDA showing the genomic variation explained by precipitation of the wettest month (bio13) and mean temperature of the wettest quarter (bio08) after correcting for neutral population structure. Circles represent populations and colors indicate the majority membership of individuals to one of three genomic clusters (cfr. The pie charts in (A)).

We cultivated samples from a minimum of 15 maternal lines for each population until the four-leaf stage, when the plant tissue was collected and lyophilized, resulting in 342 samples. We extracted DNA from these lyophilized samples on the KingFisher purification system (Thermo Fisher Scientific, Waltham, MA, USA) using a customized sbeadex DNA extraction kit (LGC, Teddington, UK). Following a modified preparation protocol of the Illumina Nextera DNA library preparation kit (Illumina, Inc., San Diego, CA, USA; Therkildsen & Palumbi, 2017), we generated libraries suitable for low-coverage sequencing. We sequenced these libraries in five lanes of an Illumina NovaSeq 6000 at Novogene (Cambridge, UK). We mapped raw reads to the *L. sativa* reference genome (2.5 GB) (annotation version 7; GCA_002870075.2; Reyes-Chin-Wo et al., 2017) using bwa-mem2 v2.2.1 (Vasimuddin et al., 2019) with default settings. Higher coverage was obtained in the genic regions (ca. 2.65X as compared to 1.76X for the entire genome), which were then retained for downstream analyses. We removed samples with average coverage < 0.5X. We used sambamba v.0.8.1 to remove low-quality mapping reads (MAQ < 20), picard tools v2.27.1 to remove PCR duplicates (*Picard Tools*, 2019), and bamutil v1.0.15 to clip overlapping reads.

### Population genetic structure

We investigated population structure using a principal component analysis (PCA) performed on the genotype likelihoods of SNPs from 800 randomly selected exons of length between 500 and 5000 bp using pcangsd v1.0 (Meisner & Albrechtsen, 2018). In our initial results, we identified eight samples from population A as outliers based on their PCA scores (Figure S4). By computing an identity-by-state relatedness matrix of SNPs using SNPRelate (Zheng et al., 2012); Figure S5), we detected that these eight individuals were close relatives while showing minimal relatedness to all other samples. This suggests that these individuals may be siblings of hybrid origin and were thus excluded from downstream analyses, thus removing their potential influence on corrections for neutral population structure. We further validated technical replicates, which were then excluded from the dataset, leaving 329 samples with a minimum of 14 individuals per population with the exception of population A (Table S1), and computed a new PCA. We then performed admixture analysis via ngsadmix for ancestries k ranging from 1 to 10 (10 replicates per k), based on genotype likelihoods calculated in angsd v0.940 (-GL 1 -doGlf 3 -doMajorMinor 1 -doMaf 2 -minMapQ 20 -minQ 20 - SNP_pval 1e-6). Results were visualized as bar plots generated by the popHelper package in R (Francis, 2017).

### Identification of climatic drivers of adaptive variation and environmental association analyses

To detect polymorphic sites (SNPs) across the exome, we ran freebayes v1.3.7 with the options of “-- min-repeat-entropy 1”, “ --max-complex-gap -1”, “--haplotype-length -1”, and “--use-best-n-alleles 4”. We calculated population allele frequencies from the inferred genotype likelihoods using the “popStats” function implemented in vcflib v1.0.3 (Garrison et al., 2022). We retained only SNPs with a minimum allele frequency (MAF) of 0.1 in at least one population and subsequently generated a folded site frequency spectrum (SFS) to remove SNPs with MAF below 0.05 for the entire dataset. This resulted in allele frequencies of 580,412 SNPs retained for downstream analyses.

We used WorldClim data for our 21 populations to explore the climatic drivers of adaptive genomic variation in a redundancy analysis (RDA). An RDA extends multiple regression to multivariate response data to assess the impact of each environmental predictor on allele frequencies across the genome. RDA was implemented using the “rda” function from the vegan package (Oksanen et al., 2022). We normalized the 19 bioclimatic variables using z-score normalization (centering and scaling) to ensure comparability and equal weighting in the RDA. To avoid multicollinearity among the environmental predictors in the RDA, we performed step-wise variable selection. We first retained precipitation of the wettest month which was correlated highly with mean annual precipitation (correlation = 0.99) and precipitation of the wettest and coldest quarters (correlation = 1.00 and 0.97, respectively), the period during which the majority of the precipitation falls. As temperature is known to be a key factor in determining the northern range limit of the species in Eurasia as a whole (Chadha & Florentine, 2021), we also retained mean temperature of the wettest quarter (bio8) to capture the temperature during which the majority of the precipitation falls (correlation with mean annual temperature = 0.83). All other variables with an absolute correlation coefficient greater than 0.8 with at least one of these two variables were removed. Among the remaining variables showing absolute pairwise correlations below 0.8, we retained those presumed to have the greatest biological importance, resulting in a final predictor variable set comprising bio2 (mean diurnal range), bio3 (isothermality), bio7 (temperature annual range), bio8 (mean temperature of the wettest quarter), bio10 (mean temperature of the warmest quarter), bio13 (precipitation of the wettest month), bio15 (precipitation seasonality), bio17 (precipitation of the driest quarter), and bio18 (precipitation of the warmest quarter).

Our RDA analysis involved comparing two models: a minimal model with an intercept only, and a full model incorporating all uncorrelated variables. A PCA performed on population allele frequencies indicated that the first three PCs explained the majority of the genomic structure, with a discrete drop in explained variance observed beyond the third component. We therefore added a condition to correct the RDA for population structure using the first three principal components. Forward stepwise model selection was then conducted using the “ordiR2step” function from the vegan package (Oksanen et al., 2022). This function systematically selected variables based on their explanatory power (*R*^2^) and statistical significance (*P* ≤ 0.06), with 100 permutations per variable to assess significance. This allowed us to determine the climatic variables explaining the largest proportion of adaptive genomic variation Although only a single precipitation variable (precipitation of the wettest month) was selected by the model as explaining a significant proportion of adaptive genomic variation (adjusted *R*^2^ 0.017; *P* = 0.032). we also retained a temperature variable, i.e., temperature of the wettest quarter, given temperature’s known biological importance to the species’ response to climate (Prince et al., 1985).

We used latent factor mixed models (LFMM) to identify SNPs that showed linear associations with precipitation of the wettest month (version 2.0; (Caye et al., 2019). This approach benefits from a low false-discovery rate while still maintaining a relatively high true-positive rate (Lotterhos, 2023). To correct for neutral population structure, we used K=3 latent factors in the models, and accounted for multiple testing by selecting SNPs with qvalues, calculated using the qvalue package in R (Storey et al., 2023), less than or equal to the False Discovery Rate of 0.05 (FDR; i.e., the expected proportion of falsely identified SNPs among all significant tests). The identified outliers formed the set of candidate SNPs occurring in genes putatively involved in adaptation to precipitation.

### Variation in selection along the precipitation gradient

We investigated the strength of selection acting on each candidate SNP across the species’ niche using modified generalized dissimilarity models (GDMs;(Dudaniec et al., 2018; Fitzpatrick & Keller, 2015; Razgour et al., 2019), using the GDM R package (Fitzpatrick et al., 2022). Originally used to assess species turnover and changes in biodiversity along environmental gradients (Ferrier & Guisan, 2006), this method applied to genomic data allows the investigation of trends in allele frequency shifts for SNPs across climatic gradients or geographic distances (Fitzpatrick & Keller, 2015). The first step in this approach is the construction of population-level pairwise genetic distance matrices for each site-by-SNP pair (Oliehoek et al., 2006). This matrix is used to predict the magnitude and rate of change in genetic distance, hereafter “allelic turnover”. I-splines are then fitted to curves of allelic turnover across the environmental gradient, with the curve’s maximum height indicating a variable’s importance in explaining turnover at each SNP. The shape of the fitted curve reveals how partial allelic turnover changes across an environmental gradient, thus highlighting locations along the environmental gradient where selection is more pronounced (Fitzpatrick & Keller, 2015).

Applying GDMs to the candidate SNPs identified through LFMM, we included precipitation of the wettest month, mean temperature of the wettest quarter, and the Euclidean geographic distance between populations as predictor variables. The inclusion of geographic distance acted as an additional test to determine if neutral processes, such as isolation by distance, rather than isolation by environment, drive turnover. Temperature of the wettest quarter was included to identify SNPs with a stronger response to temperature than to precipitation. We generated pairwise matrices of genetic distances between each of the 21 populations independently for each of the outlier SNPs using the R package HIERFSTAT (Goudet, 2005; Nei, 1987). We then retained a single SNP per gene, selecting those SNPs with the greatest deviance in allele turnover explained by the model, because the fit of a GDM is assessed by the percentage of deviance explained by the model (Ferrier & Guisan, 2006; Fitzpatrick & Keller, 2015; Fitzpatrick et al., 2013). This resulted in 230 SNPs, of which 221 had converged models and were retained for further analysis. We then calculated the relative importance of each environmental variable for each SNP by normalizing the maximum height of the I-spline curve (i.e., cumulative turnover in allele frequency) to between 0-1 using min-max normalization, generating a variable called “relative importance”.

We used GDM results to further filter our candidate SNPs for potential false positives, despite the implementation of latent factors in our LFMM. We retained SNPs with a strong association with precipitation of the wettest month (relative importance >0.95), a weak association with geography (<0.05), and relative importance of precipitation higher than for temperature of the wettest quarter.

After applying these filters, 98 SNPs remained and their relative importance was newly rescaled. We classified SNPs into three groups based on the region along the precipitation of the wettest month gradient in which they showed the steepest allelic turnover. Specifically, SNPs were classified as those with the steepest turnover at the dry niche edge (50-100 mm; hereafter “niche edge SNPs”; Figure S2), in the middle of the gradient (100-150 mm, hereafter “intermediate SNPs”), and at the mesic end of the gradient (150 - 200 mm; hereafter “niche center SNPs”).

### Genetic basis of adaptation to precipitation

We identified functions of genes associated with the candidate SNPs using the annotation of the *L. sativa* genome. We used the GO terms of these *L. sativa* genes to perform enrichment analyses of biological functions for the entire set of filtered SNPs resulting from the GDM analysis, as well as for the separate GDM groups at the niche edge and niche center. To ensure a conservative test, we required a minimum of 4 nodes and used the Fisher’s Exact Test with the ‘elim’ algorithm. Only GO processes with *P*<0.005 were retained. We further assessed the function of genes included in enriched categories through their available annotation in their respective *Arabidopsis thaliana* orthologs.

### Adaptive genetic composition along the precipitation gradient

While GDM allows us to investigate patterns of selection at the level of individual SNPs, it does not allow us to examine the polygenic signal of adaptation across populations. We explored how the cumulative adaptive genetic composition of populations varies along the precipitation gradient to assess potential shifts in selection regimes across the niche of the species. To do this, we estimated polygenic risk scores (PRS) for each population, a method originally developed in medicine to estimate individual risk for diseases based on genetic markers and their strength of association with a condition (Dudbridge, 2013; Wray et al., 2007). PRS was calculated as the cumulative sum of the allele frequencies for the precipitation-associated SNPs for each population, with each SNP weighted by its effect size from the LFMM. In practice, this method integrates the contribution of each SNP to generate the polygenic composition of each population as a result of adaptation to the precipitation regime experienced in their respective habitat. A consistent selection coefficient along the precipitation gradient, paired with adaptation, is expected to result in linear changes of allele frequencies, and ultimately a high correlation between the PRS values and precipitation. Conversely, variation in selection pressures along the gradient could lead to a non-linear PRS trend.

To ensure unbiased PRS estimates, we filtered the retained set of 98 SNPs for linkage disequilibrium, calculated using the GUS-LD package in R (Bilton et al., 2018). For each pair of SNPs showing *r*^2^ > 0.8 and occurring on the same chromosome, we retained the SNPs with the best model fit under the GDM, resulting in a dataset of 59 unlinked SNPs (Figure S6). We fitted both linear and non-linear models with precipitation as the response variable and PRS as the explanatory variable to the data and compared their model fit and explanatory power. We further compared the best-fitting curve to the expected null relationship between precipitation and PRS based on sets of 59 randomly chosen SNPs and their effect sizes from the LFMM model. For each of 1000 replicate runs of 59 randomly chosen SNPs, we extracted fitted values from linear models between precipitation and PRS values, and displayed the expected relationship as the 2.5th and 97.5th percentiles of the fitted values.

### Temporal projections of adaptive evolution in response to climate change

To explore putative evolutionary responses of local populations to shifts in precipitation regimes linked to climate change, we projected the function describing the association between the adaptive genetic composition of populations and precipitation, i.e., our PRS curve, to both past and future climatic predictions across Israel. As there has been an increase in the length of the summer drought period in Israel between 1975 and 2020 (Drori et al., 2021), we contrasted the PRS model as applied to precipitation of the wettest month values for the periods of 1965 to 1974 and 2010 to 2019. We downloaded monthly precipitation data at a 2.5 arc-minute resolution from WorldClim for these periods and applied the model to generate a climate raster (Fick & Hijmans, 2017), enabling us to project PRS for *L. serriola* populations across Israel. We forecast these PRS patterns into the future using the MIROC 6 global climate model (GCM) (Tatebe et al., 2019) generated as an estimate for the period 2061-2080 under the most extreme scenario of SSP585 with downscaled data made available through the Coupled Model Intercomparison Project Phase 6 (CMIP 6) and accessed through WorldClim (Eyring et al., 2016). By comparing PRS values between these intervals, we identified putative spatial trends of adaptive evolution over time.

## Results

### Neutral population structure of *L. serriola* in Israel

Populations of *L. serriola* were composed of a mixture of individuals with varying ancestry, with individuals sometimes clustering more closely with other populations than with the populations from which they were sampled (Fig 1B). Populations in the north (populations K, L, O-S) hosted the largest proportion of genetic variation, with individuals aligning along PC1 of the PCA (26.7% of explained variation), while variation among the wettest populations (Q-U) was mainly explained by PC2 (6.1%). Individuals from southern, drier populations were differentiated primarily along PC3 (4.7%). Thus, the first three principal components explained a large proportion of genetic differentiation along the geographic cline (37.5%), and similar results were obtained when the PCA was computed on population allele frequencies (65%; Fig S8). In both analyses, the eigenvalue plots indicated a discrete drop in explanatory values after the first three principal components (Figure S7 and S8). Genetic admixture computed for K=3 followed a latitudinal pattern, with southern, central, and northern populations forming relatively distinct groups (Figure 1A, Figure S9).

### Adaptive genetic variation along the precipitation gradient

Precipitation (bio13) was the only significant driver of adaptive genetic variation retained by the RDA during forward model selection. Neutral structure explained 73.6% of the genomic variation, 4.5% was explained by environmental variables, and 21.9% remained unexplained. After correcting for neutral structure, RDA1, which was predominantly correlated with precipitation of the wettest month, captured 11.24% of the remaining variance and RDA2, more strongly correlated with mean temperature of the wettest quarter, 5.81% (Figure 1C). LFMM performed on precipitation of the wettest month identified 488 significantly associated SNPs (Figure 2A; Figure S10) occurring in 230 genes across the nine chromosomes.

**Figure 2:**
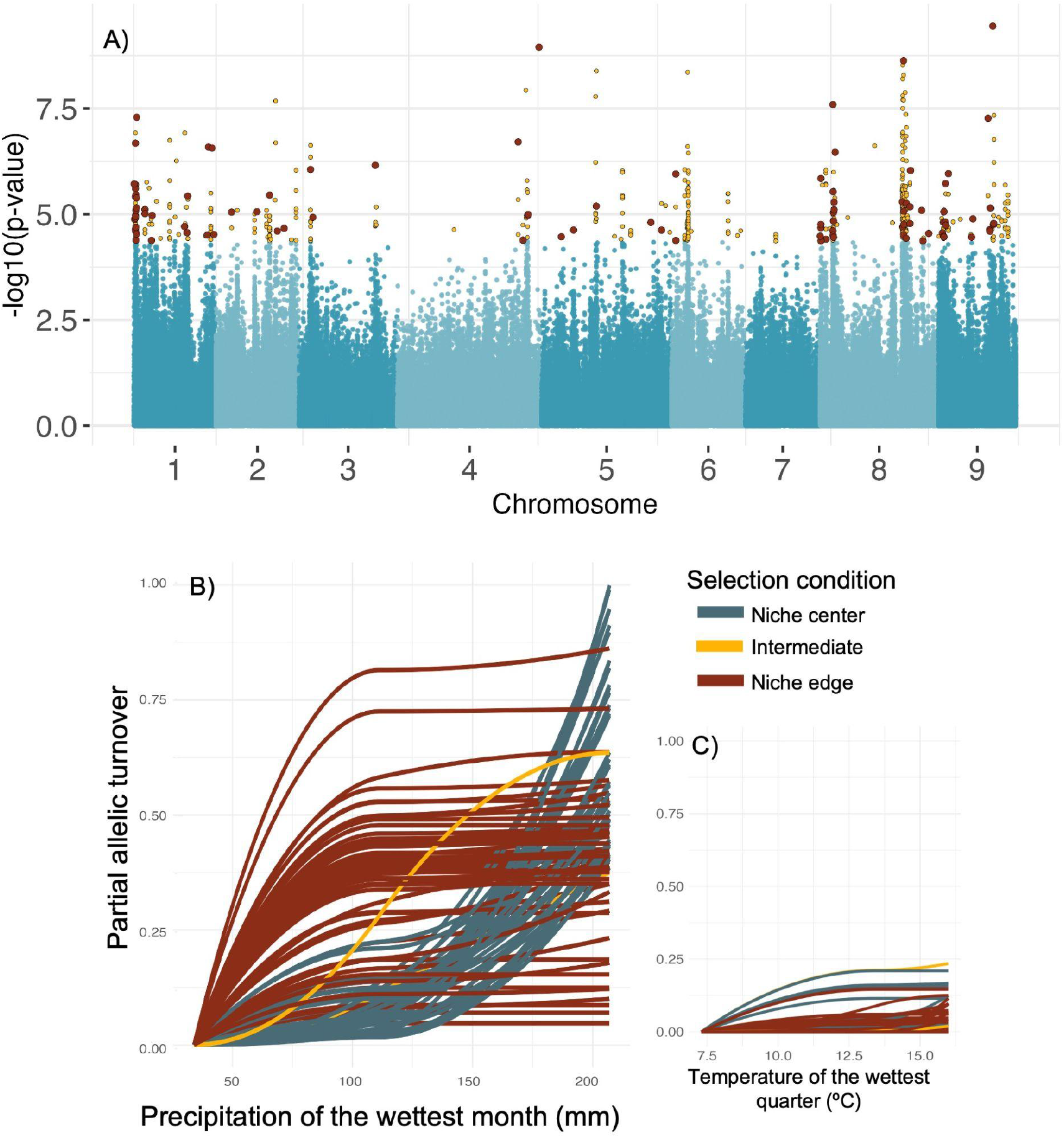
Adaptive variation along the precipitation gradient in Israel. (A) Environmental associations of genomic variation with precipitation identified using a latent factor mixed model (LFMM). Light and dark blue points denote SNPs on alternate chromosomes, and yellow points denote SNPs significantly associated with precipitation of the wettest month (n=488). SNPs retained after filtering for potential false positives associated strongly with geography and temperature in the GDM analyses are shown in red (n=98). (B) Partial allelic turnover of candidate SNPs. SNPs are classified according to their highest turnover at the arid niche edge (50-100 mm; red), intermediate (100-150 mm, yellow), and the transition to more mesic conditions (150-200 mm, blue). (C) Turnover for these same candidate SNPs visualized across temperature.

To identify variation in the strength of selection acting on these 488 SNPs, we computed partial allelic turnover using GDM analyses implementing geography, precipitation of the wettest month, and mean temperature of the wettest quarter as predictors, and retaining a single SNP per gene resulting in a dataset of 221 SNPs that achieved model convergence. Of these SNPs, 111 showed relatively strong associations with geography, and 13 showed stronger associations with temperature than with precipitation. Removing these latter SNPs resulted in a dataset of 98 candidate SNPs.

Allelic turnover for the 98 candidate SNPs did not follow a linear pattern along the precipitation gradient (Figure 2B). Specifically, 54 SNPs exhibited highest turnover near the arid niche edge, while 42 SNPs exhibited the highest turnover near the more mesic niche center, and two under intermediate conditions. These SNPs showed minimal turnover with temperature (Figure 2C). After removing highly linked SNPs, we observed a larger difference in the number of SNPs under selection at the niche edge compared to the center, with 42 exhibiting the highest turnover at the niche edge, and 15 exhibiting the highest turnover near the niche center (Figure S11).

### Genetic basis of adaptation to precipitation

Enrichment analyses conducted on the set of 98 candidate genes revealed several significantly enriched GO terms in line with known biological processes involved in plant response to selection imposed by precipitation. These included GO terms related to water deprivation, and defense from viruses, bacteria, and other organisms. Interestingly, separate analyses of SNPs under selection at the dry niche edge and nearing the niche center revealed almost unique sets of enriched biological functions. Niche edge SNPs showed enrichment of regulation of protein deubiquitination (LATERAL ROOT STIMULATOR 1 [LRS1]), formaldehyde catabolic process (AT5G42250; AT5G42250), and TOR signaling (LETHAL WITH SEC THIRTEEN 8-1 [LST8-1]) (Figure 3b). Further, we also identified genes involved in biological processes related to photoperiodism and flowering (MORF RELATED GENE 1 [MRG1]; WITH NO LYSINE KINASE 5 [WNK5]; PWWP DOMAIN PROTEIN 1 [PDP1]; AT1G18560; CENTER CITY [CCT]; LETHAL WITH SEC THIRTEEN 8-1 [LST8-1]), regulation of reproductive process (TETRATRICOPEPTIDE REPEAT 2 [TPR2]), and cellular or organismal response to starvation of nutrients or energy (HOMOLOG OF YEAST AUTOPHAGY 18 [ATG18] H [ATG18H]). Niche edge SNPs also showed enrichment in general defense, as well as defense responses to symbionts, viruses, and fungi.

**Figure 3:**
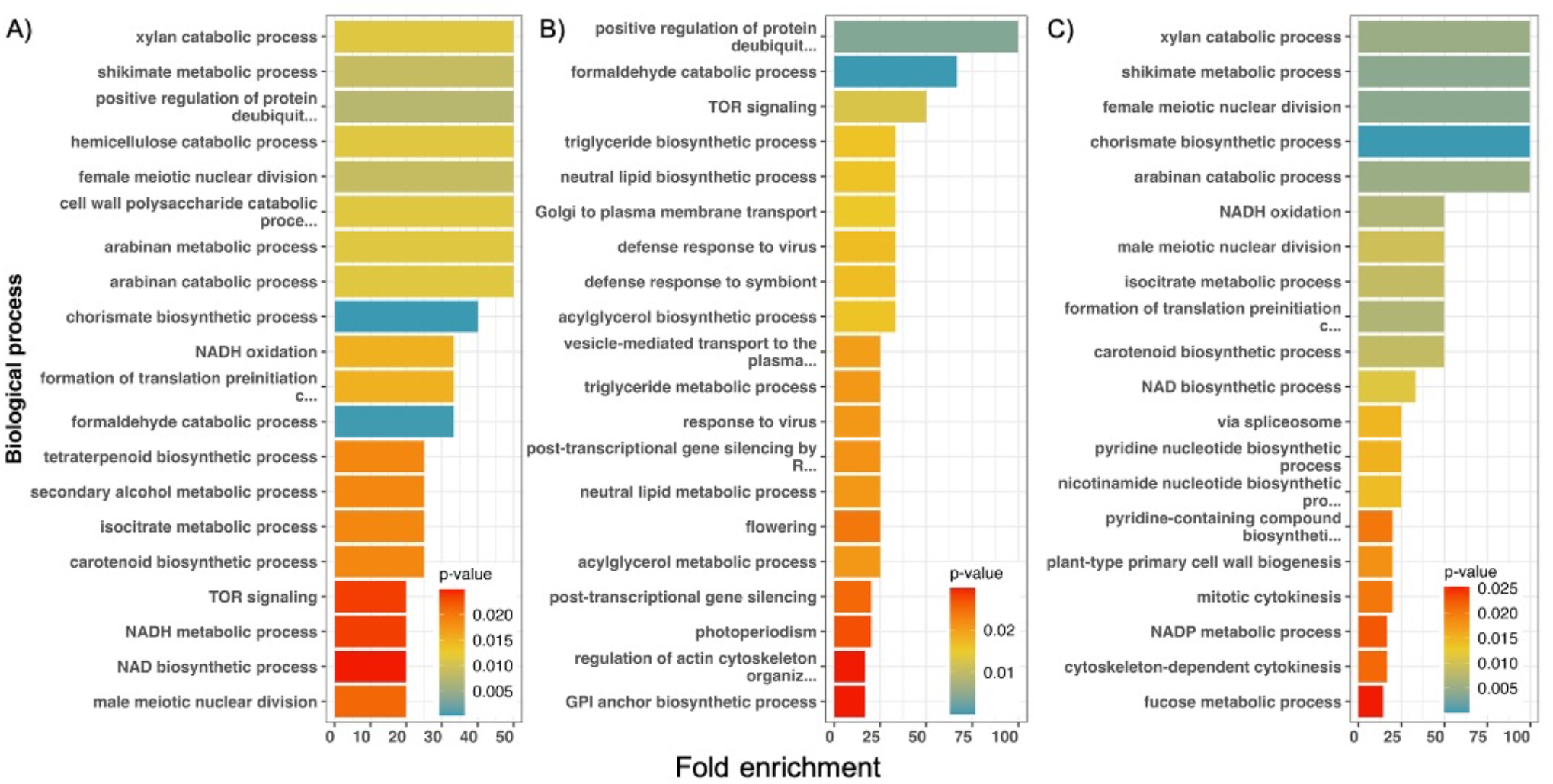
Fold enrichment for biological processes of genes containing candidate SNPs. Significance from the Fisher exact test indicates whether the proportion of genes with a specific GO term in the gene set of interest deviates from what would be expected based on their prevalence in the reference genome (colors). Fold enrichment indicates the relative increase in the number of genes with each GO term compared to what would be expected randomly with the same sample size from the genome. Depicted are the fold enrichments for GDM-filtered SNPs (A; n=98), niche edge SNPs (B; n=54), and niche center SNPs (C; n=42). Only the top twenty most enriched GO terms are shown. All GO terms have *P*-values below 0.05.

SNPs under selection near the more mesic niche center showed significant enrichment in xylan catabolic processes (AT1G78060), shikimate metabolic processes (MODIFYING WALL LIGNIN-1 [MWL1]), as well as both female and male meiotic nuclear cell division (ZYP1a; X-RAY INDUCED TRANSCRIPT 1 [XRI1]). A single phenology gene was also identified as possibly under selection near the niche center (EARLY BOLTING IN SHORT DAYS [EBS]), as well as a gene related to flower development (AGAMOUS-LIKE 8 [AGL8]) and another to the regulation of reproductive processes (HISTONE 3.3 [H3.3]). Several genes related to plant-type cell wall biogenesis were found to be enriched in the niche center SNP dataset (AT3G01190; AT1G78060; CELLULOSE SYNTHASE 6 [CESA6]). Both niche center and edge SNP datasets showed enrichment in genes related to general defense response; however, only more mesic niche center SNPs were enriched in defense responses to bacteria and to other organisms.

### Adaptive genetic compositions and temporal projections of putative evolutionary change

Consistent with patterns of allelic turnover found in the GDM analyses, PRS computed on the dataset of 59 unlinked SNPs was best fitted by a cubic regression between adaptive differentiation (PRS score) and precipitation of the wettest month (estimate ± error: 5.81e-08 ± 1.96e-08; p<0.001; Table S3). Specifically, we found a steep curve reflecting strong adaptation near the niche edge, while populations above 100 mm, closer to the niche center, showed relatively weak adaptive differentiation.

Our temporal projects suggest that arid-associated alleles have increased in frequency throughout the range since 1975, especially in the north of Israel where the climate has become increasingly dry. By contrast, adaptive genetic composition remained relatively more stable in central Israel during this same period. Future projections show that even under relatively strong climate change scenarios, shifts in allele frequencies will be marginal compared to those that might have occurred in the past. These are projected to affect primarily populations in regions with mesic precipitation regimes in the north and in populations falling around the Sea of Galilee and neighboring the Jordanian Highlands (Figure 5e). Conversely, the arid southern populations and more central populations under moderate precipitation regimes were predicted to show little to no change in adaptive genetic compositions. Along the southern coast, populations are predicted to shift to wetter genetic compositions in response to a local change in climate towards more mesic conditions.

**Figure 4:**
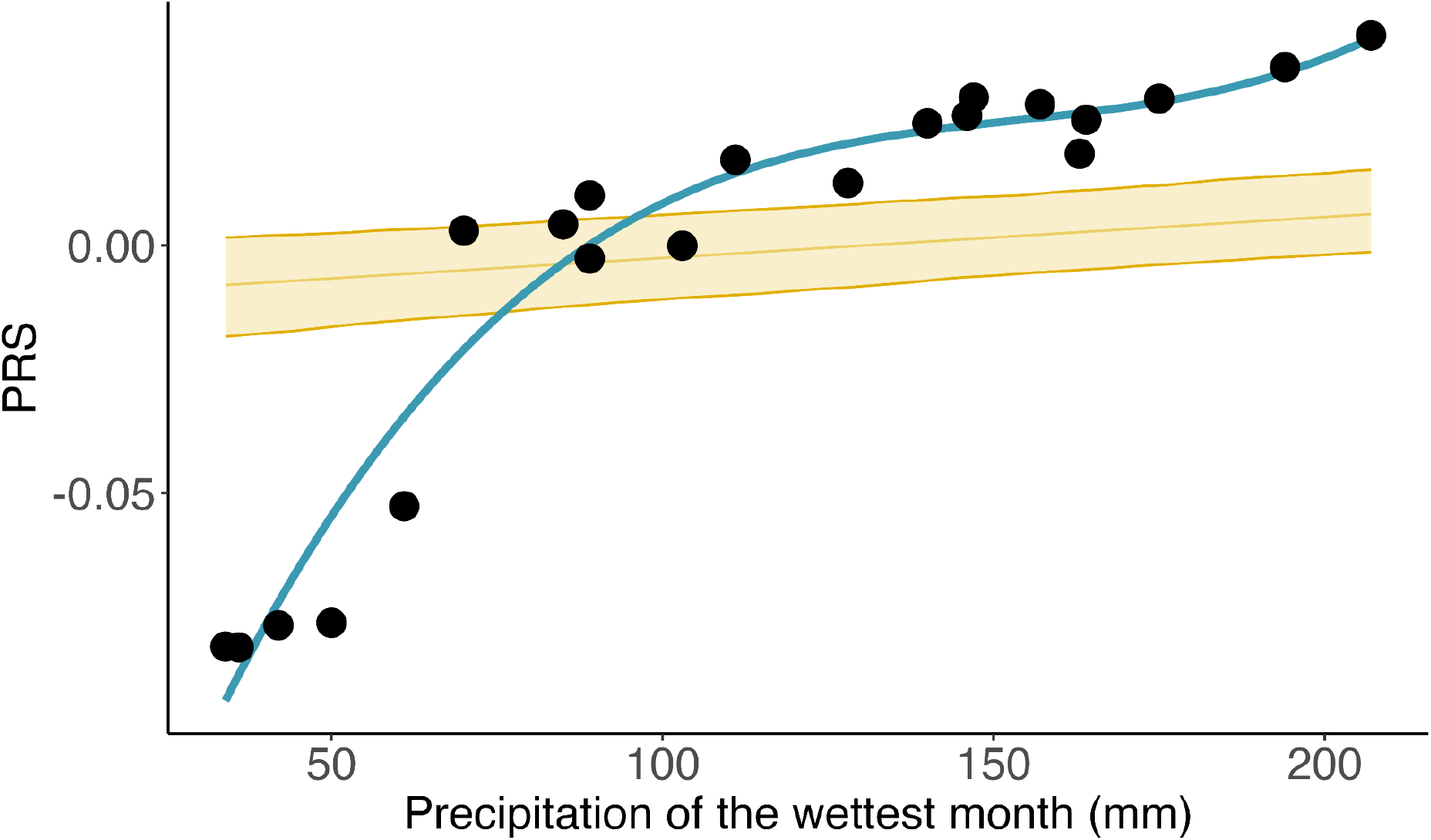
Polygenic risk scores (PRS) of 21 populations of *L. serriola* across the precipitation gradient in Israel. Populations PRS scores (points) fitted with a sigmoid curve for the data set of unlinked SNPs (n=59). The null model, depicted with the 95% confidence interval (yellow), is generated for the same number of randomly selected neutral SNPs bootstrapped for 1000 iterations.

**Figure 5:**
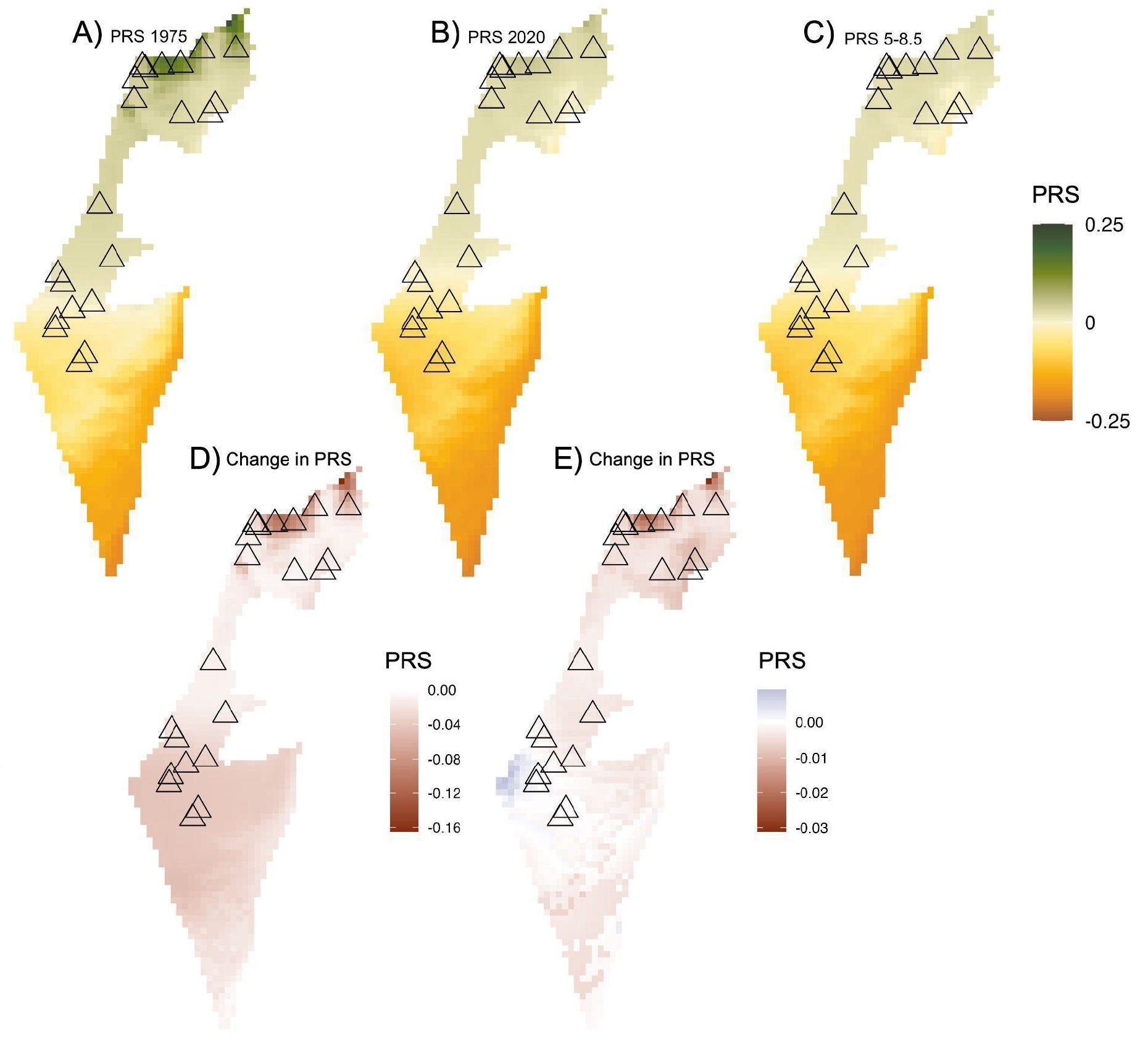
Projected changes in genetic composition of adaptive SNPs in response to shifts in precipitation regimes as a consequence of climate change. Predicted PRS for (A) 1975, (B) 2020, and (C) 2070 under climate scenario SSP5-8.5 (highest emissions) for precipitation of the wettest month. Triangles indicate the locations of populations included in the study. (D, E) The change in PRS scores indicates the predicted evolutionary change between the two time periods above (A and B, or B and C; respectively). Positive values indicate a shift in allele frequencies corresponding to more mesic conditions, while negative values suggest shifts to drier genetic compositions. Note that the scales of the “change in PRS” maps vary compared to that of the year-specific PRS maps and to each other.

## Discussion

### Climate adaptation in a ruderal species

Israel’s steep precipitation gradient serves as an ideal backdrop to examine potential adaptive responses of plant species to changing climate over space and time. We examined how the strength of selection and adaptation to precipitation varies from the center to the edge of the climatic niche of *Lactuca serriola,* and identified genes and corresponding functions underlying adaptation along this precipitation gradient. *L. serriola* is a ruderal species with a high dispersal and selfing rate (Chadha & Florentine, 2021; D’Andrea et al., 2016; Mejías, 1994; Weaver & Downs, 2003), implying that similar genotypes carried by full-siblings are likely to disperse into areas that experience different selection regimes across the species’ distribution. Consistently, individuals from the same population did not necessarily share similar ancestry and showed relatively low relatedness. This suggests that there is a high degree of gene flow between nearby populations, and also over relatively long distances across the geographic range. High mobility, paired with potential anthropogenic dispersal, could in principle impede the ability of populations to adapt locally if gene flow is higher than the strength of selection (Bridle & Vines, 2007; Tigano & Friesen, 2016). However, we found that precipitation was a strong driver of adaptive genomic variation, indicating that selection associated with precipitation has been strong enough to overcome the homogenizing effect of dispersal. Nonetheless, the nonlinear association between the adaptive genetic composition of populations and local precipitation regimes indicates that the strength of selection is not even along the gradient.

### Variation in selection across Israel’s precipitation gradient

The majority of associated and unlinked SNPs were found to be under stronger selection near the dry niche edge, while a much smaller number were found under the more mesic conditions near the niche center. Theory predicts selection to be weaker or stabilizing near the center of the niche, and to increase and become more directional closer to the niche edge (Angert et al., 2020; Brown, 1984; Hargreaves et al., 2014; Lee-Yaw et al., 2016). Our results are in line with such predictions, since populations close to the niche center showed more similar adaptive genetic composition (PRS scores) in contrast to populations occurring toward the niche edge, which displayed relatively strong divergence in genetic compositions. Interestingly, SNP-specific patterns of allelic turnover suggest that different sets of SNPs may contribute to adaptation in the dry and mesic portions of the climatic niche. Biological processes for genes containing candidate SNPs with the strongest partial allelic turnover near the niche edge are enriched with TOR signaling, which helps regulate plant response to osmotic stress (Fu et al., 2020). Moreover, enriched terms include flowering related processes and deubiquitination, also known to influence flowering time (Schmitz et al., 2008). Phenology is highly responsive to changes in climate (Anderson et al., 2012; Forrest & Miller-Rushing, 2010), and phenological shifts in response to climate have been observed in *L. serriola* (Alexander, 2013; Prince et al., 1978). Earlier phenology is a key trait often associated with drought avoidance, (Kigel et al., 2011), which together with shorter lifespans and rapid growth, enables plants to complete reproduction before the onset of drought conditions (Kooyers, 2015).

Biological processes of genes containing SNPs with the strongest partial allelic turnover near the niche center showed enrichment in defense against other organisms, xylan catabolic process, and cell-wall biosynthesis. Both the xylan catabolic process and cell-wall biosynthesis can be important in preventing cellular damage from fungal infection (Beliën et al., 2006; Wan et al., 2021). While *Lactuca* pathogens have varying precipitation requirements (Mieslerová et al., 2020), and there are a few plant-pathogen interactions that are more severe under drought conditions, most plant-pathogen interactions become more virulent with increasing humidity and rainfall (Velásquez et al., 2018).

Furthermore, the enrichment of biological processes related to defense against other organisms is consistent with biotic interactions becoming more important under more benign environments, as formulated by the Species Interactions–Abiotic Stress Hypothesis (Louthan et al., 2015; Paquette & Hargreaves, 2021). If biotic interactions do indeed impose stronger selection under more benign conditions towards the niche center, the potential polygenic architecture of adaptation involving many genes of small effect that may be particularly difficult to detect with EAA methods (Rellstab et al., 2015; Wellenreuther & Hansson, 2016). This might also explain why we observed a flattening of the relationship between PRS and precipitation in more mesic environments. While future work is needed to validate that these functions differ along the precipitation gradient, our findings are consistent with previous studies which suggest that biotic interactions are stronger under mesic conditions, while increasingly arid conditions impose selection for stress tolerance (Liancourt & Tielbörger, 2009; Schiffers & Tielbörger, 2006).

### Implications for range limits and climate adaptation

Although *L. serriola* occupies very arid environments at its range edge in Israel, drier climates exist in Israel which remain uninhabited (Figure S3). This raises the question as to what determines the observed dry limit of the species’ range. Firstly, the arid niche edge in Israel is the Arava Desert, which may not support the biotic or non-climatic abiotic conditions needed for the species to establish. Beneficial below-ground symbionts, facilitative biotic interactions, or appropriate soil conditions may be lacking, preventing establishment in drier conditions. This seems unlikely, however, as the species is ruderal and found to establish in various soil conditions (Chadha & Florentine, 2021), and the growth and competitive ability of *L. serriola* is often negatively affected by the bacterial communities in the rhizosphere (Aguilera et al., 2017). Alternatively, gene flow between neighboring populations close to the niche edge may prevent adaptation to more arid conditions (Kirkpatrick & Barton, 1997), especially along the steepening environmental gradient that we observed (Bridle et al., 2019). If range edge populations are small, gene flow from neighboring populations under slightly more mesic conditions may have a particularly strong effect on genetic composition of edge populations, preventing further adaptation (Bridle et al., 2019; Eckert et al., 2008). However, populations in the south were not substantially smaller than those closer to the niche center (Figure S12). Given this, the most likely explanation for the lack of adaptation to even drier available climates and the formation of the niche edge is a lack of adequate genetic variation. This could be in terms of single traits for which variation is exhausted by selection at the range edge, or perhaps more plausibly by a lack of multivariate trait variation caused by antagonistic fitness trade-offs between traits required for evolution and adaptation to drier climates (Blows & Hoffmann, 2005; Bridle & Vines, 2007; Etterson & Shaw, 2001; Sexton et al., 2009). For example, trade-offs between the size and timing of reproduction have been implicated in setting range limits in other species (Bridle & Vines, 2007; Colautti & Barrett, 2013).

The climate in Israel has changed substantially over the past 45 years, particularly through an extension of the summer drought period (Drori et al., 2021). In conjunction with the steep precipitation gradient across Israel, *L. serriola* offers the opportunity to consider how populations may have shifted their adaptive genetic composition in response to these changes in precipitation regimes. Predicted spatial changes in the genetic composition of populations between 1975 and 2020 suggest that *L. serriola* has evolved in response to shifts in selection imposed by precipitation across this time. This rests on the assumption implicit in EAA methodology that populations are adapted to their current conditions, and that the genetic basis of adaptation is conserved across time. Alternatively, our signal may result from populations in the mesic range being maladapted to climatic conditions in 1975, though this seems unlikely given that mesic conditions constitute the ancestral part of the niche, and given *L. serriola’s* ability for rapid adaptation (Alexander, 2013). Potential evolutionary responses over the last 45 years would have been particularly pronounced in the north of Israel, which experienced large decreases in precipitation (Figure S13D). In contrast, populations near the dry niche edge have experienced a much smaller absolute shift in precipitation in what has always been a very dry habitat

Future projections suggest that further evolutionary change will occur primarily in northern populations, where greater changes in precipitation are expected (Figure S13E). However, when compared with the past, future adaptive shifts are predicted to be relatively small. This is partly due to the smaller predicted decrease in precipitation than has occurred in the past, but also the relatively weaker selection acting on populations in the more mesic, northern portion of the species’ niche. If conditions in the north were to dry further, populations close to the inflection point of the PRS curve (Figure 4) will face a sudden shift in selection regimes. Arid populations are predicted to have little to no shift in allele frequencies due to the very limited change in precipitation over this period. Together, these predictions of both past and future adaptive shifts illustrate that the adaptive response under warming will depend on two things, both of which can change spatially: 1) the absolute change in climate itself, and 2) the underlying set of adaptive alleles and their interaction with these climatic shifts, which leads to heterogenous selection regimes. For *L. serriola*, this results in the somewhat counter-intuitive conclusion that future adaptive shifts will be strongest towards, but not at, the niche center, where selection is weaker but precipitation changes are relatively large. In contrast, despite evidence for strong selection at the niche edge, adaptive shifts are not expected here because projected climate changes are modest. This can be explained by the strongly arid climate at *L. serriola’s* niche edge, where opportunities for further reduction in precipitation are limited; for species with range limits occurring in regions with more pronounced climate change, the necessary adaptive responses might be greater. However, the effective realization of such responses will depend on the populations’ ability to achieve novel adaptation to a drier climate leveraging on the available substrate of genetic variation, effectively expanding the current species’ niche (Moran & Alexander, 2014).

## Conclusions

Our study demonstrates that species’ responses to climatic selection can be non-linear across an environmental gradient, with an increased strength of selection towards the niche edge. These findings suggest that the magnitude of change in adaptive genetic composition in response to future climate change will depend on where populations fall within the species’ climatic niche, and the presence of standing variation required to adapt. While we highlight potentially adaptive loci and their resulting gene functions in association with precipitation, their effective role in biological processes remains to be validated. Common garden studies are needed to disentangle if there are indeed adaptive phenotypic differences in phenology, drought tolerance, or pathogen resistance, competitive ability, or other traits identified in the genes under selection along this precipitation gradient. For *L. serriola*, and other species of commercial interest for which germplasm collections exist, the evolutionary shifts we have inferred might be validated experimentally. As *L. serriola* is an agricultural weed and the wild ancestor of *L. sativa*, seed banks have accessions from around the world, some dating back several decades, which could be used in “resurrection” studies to directly test the ability of populations to adapt in a time-for-time study (Etterson et al., 2016; Franks et al., 2018; Franks et al., 2007). In sum, our study highlights the potential importance of spatial variation in selection and adaptation, its consequences for potential evolutionary responses, and the need to consider non-linear patterns of selection across spatial scales when predicting future responses of species to climatic change.

## Acknowledgments

We thank Alex Widmer for essential support with lab resources, Claudia Michel and Patrick Ackermann for molecular work, and the Genetic Diversity Centre at ETH Zurich, particularly Niklaus Zemp for providing bioinformatics support. We also thank Tomer Faraj and the seed collection team for field work. This work was supported by ETH Research Grant ETH-29 19-2.

