## Supplemental material for "Adaptation across a precipitation gradient from niche center to niche edge"

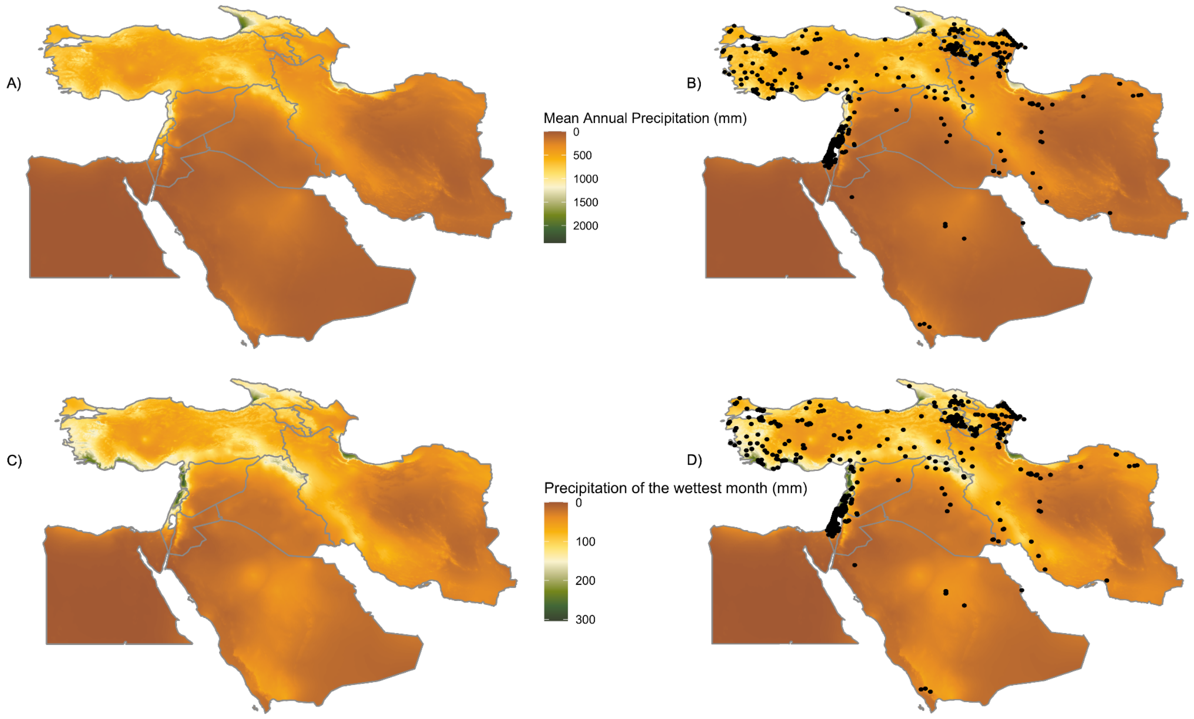


**Figure S1: Precipitation maps and occurrence of *L. serriola* in the geographic area around Israel.** (A) Climate data were extracted from 2.5-arcminute WorldClim bioclimatic variables (Fick & Hijmans, 2017). Species' occurrences (black points) were downloaded from GBIF on 16.09.2022 and cleaned using the CoordinateCleaner package in R (GBIF.org, 16 September 2022; Zizka et al., 2019). Maps with (B, D) and without (A, C) occurrence points are included to allow visualization of the underlying climate.

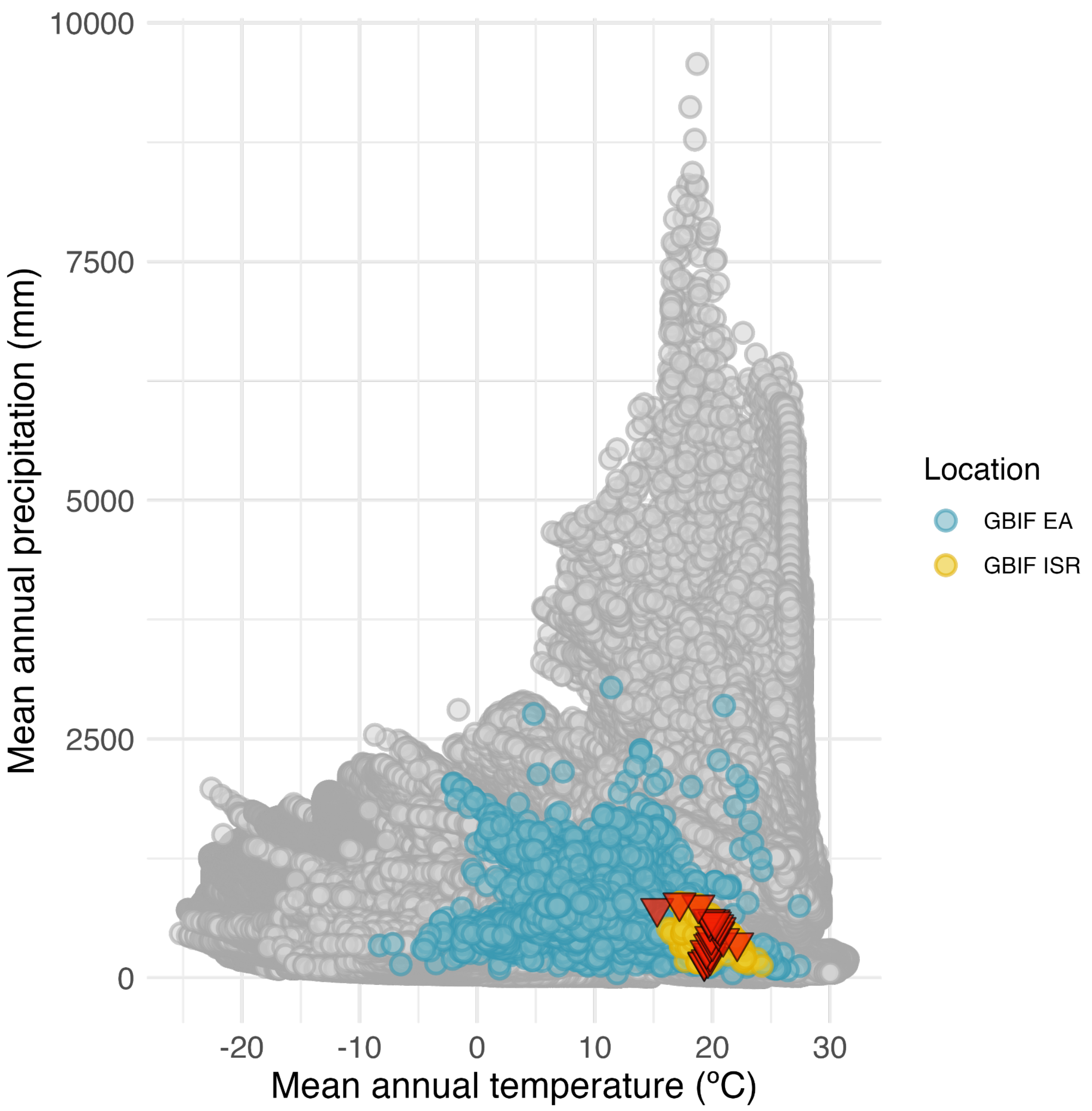


**Figure S2: *L. serriola* climatic niche for the Eurasian range of the species.** The available climate of Eurasia was estimated by extracting climatic data for all of Eurasia (n=4,299,655 points; grey points), GBIF occurrences from Eurasia (n=100,867; blue), and occurrences in Israel (n=586; yellow). The climatic locations of populations sampled (n=21) are indicated by red triangles. Occurrences were downloaded from GBIF on 16.09.2022 and cleaned using the CoordinateCleaner package in R (GBIF.org, 16 September 2022; Zizka et al., 2019). Climate data were downloaded from WorldClim at 2.5-arcminutes and includes all points for the Eurasian continent (Fick & Hijmans, 2017).

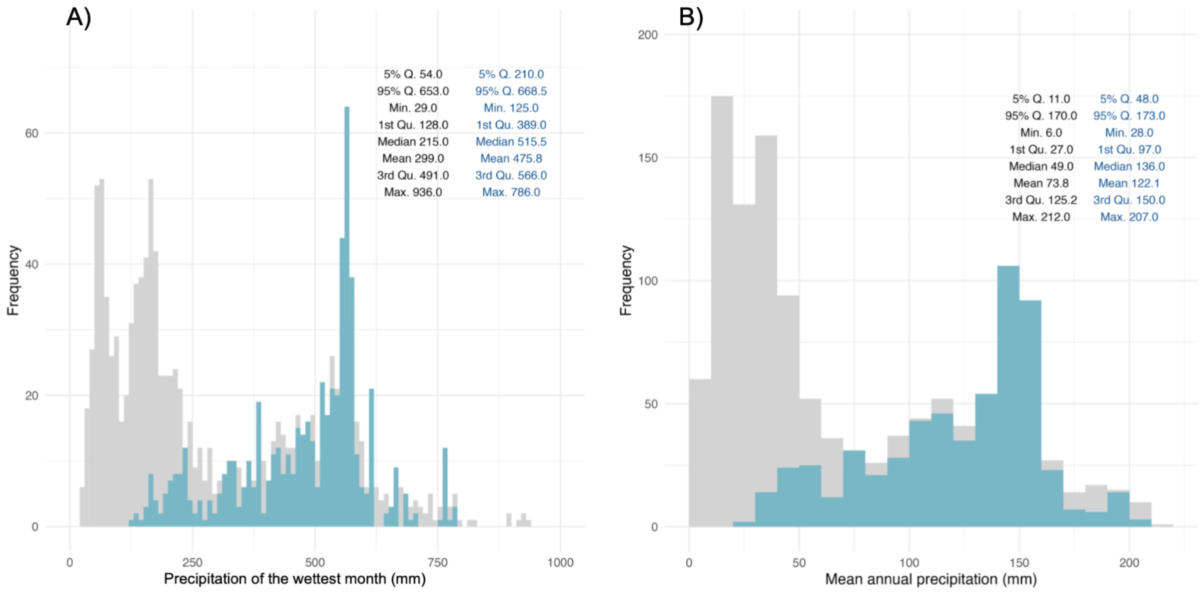


**Figure S3: Frequency of *L. serriola* occurrence along the precipitation gradient of Israel.**

Frequency of the available climate for the entirety of Israel (grey) and the occupied climate of *Lactuca serriola* in Israel (blue) with statistical summary data for the background and occurrence data (black and blue, respectively). The 5% and 95% quantiles are indicated by "5% Q." and "95% Q.". Occurrences were downloaded from GBIF on 16.09.2022 and cleaned using the CoordinateCleaner package in R (GBIF.org, 16 September 2022; Zizka et al., 2019). Climate data were downloaded from WorldClim at 2.5-arcminutes and includes all points for the Eurasian continent (Fick & Hijmans, 2017).

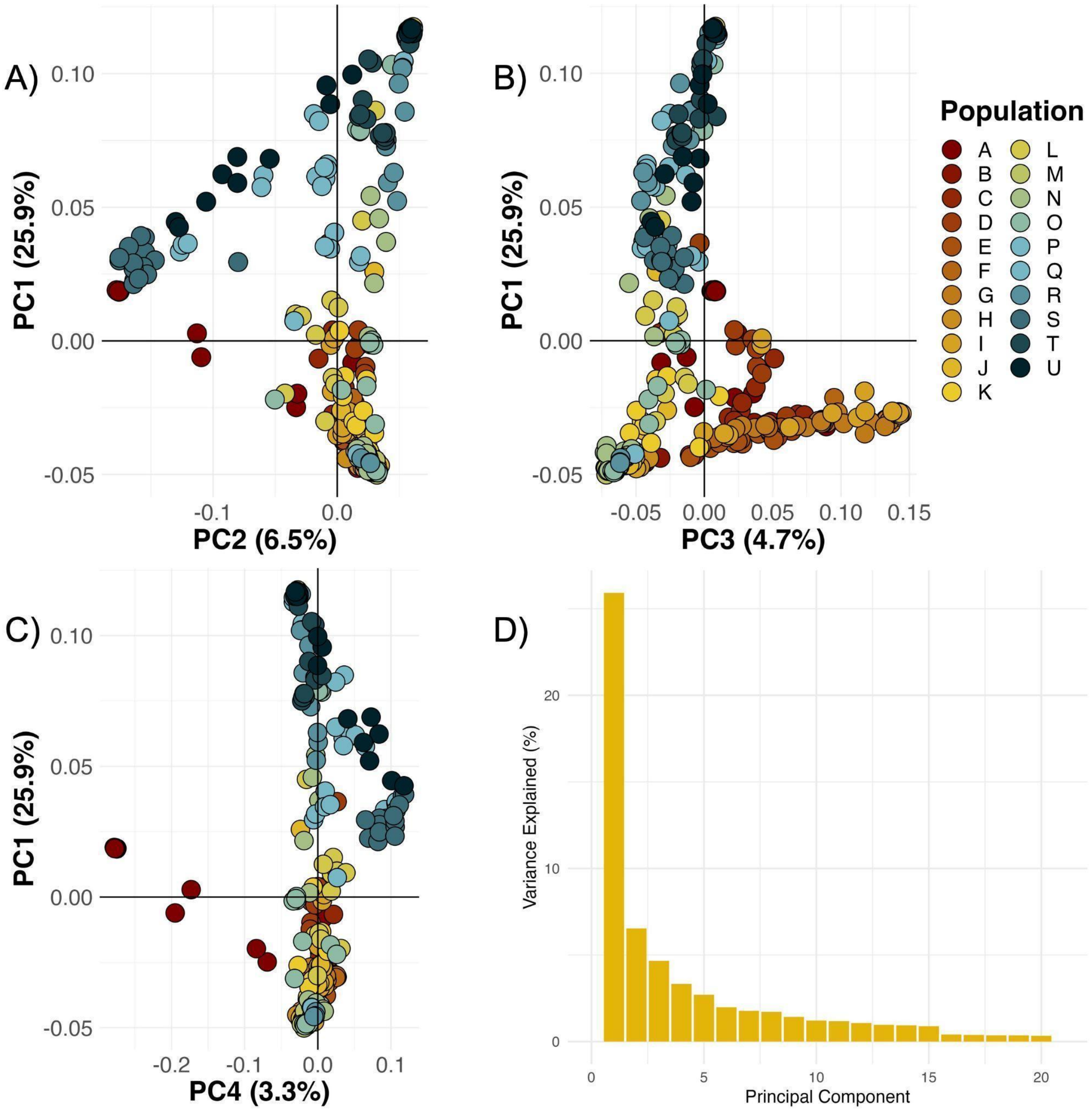


**Figure S4: PCA of individual genotypes included in the original sampling.** (A-C) PCA of genotype likelihoods for all samples (n=338) and (D) the variance explained by the principal components of the PCA. The cluster of eight samples with the lowest values on PC4 belonging to population A were removed from further analyses.

**Table S1: Sample number per population after removing eight samples of population A.**

| A | B | C | D | E | F | G | H | I | J | K | L | M | N | O | P | Q | R | S | T | U |
|---|---|---|---|---|---|---|---|---|---|---|---|---|---|---|---|---|---|---|---|---|
| 7 | 15 | 15 | 15 | 15 | 14 | 20 | 18 | 18 | 15 | 14 | 14 | 18 | 17 | 17 | 15 | 19 | 19 | 15 | 14 | 15 |

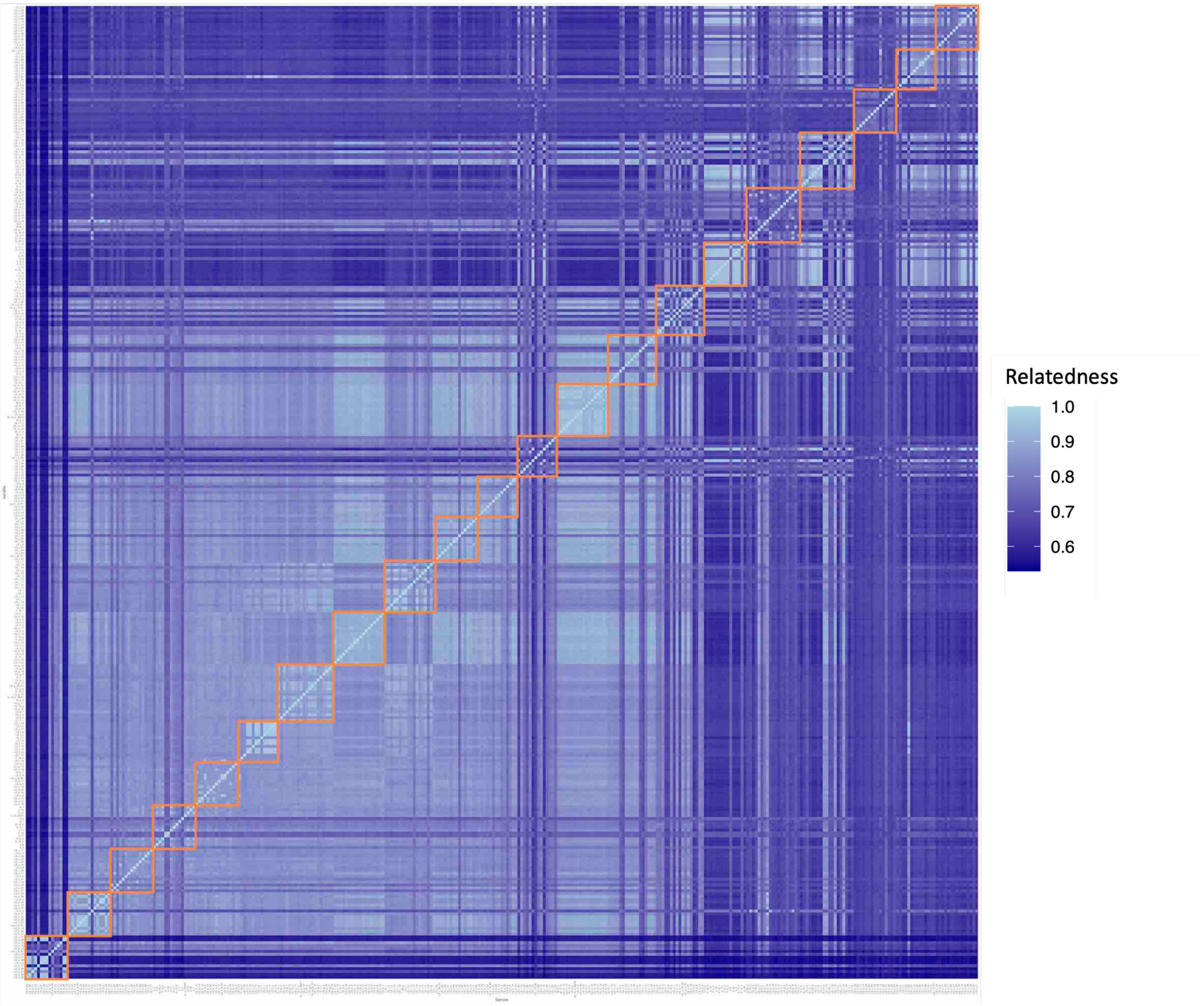


**Figure S5- Relatedness of individual genotypes included in the original dataset.** Relatedness (identity-by-state) was calculated from genotypes using SNPRelate (Zheng et al., 2012). High relatedness is indicated by lighter colors, while darker colors indicate lower relatedness between samples. Samples within the same population are highlighted in orange. The eight samples that were removed for subsequent analysis can be seen in the bottom left square, which shows high relatedness to other samples in the populations, and low relatedness to any other individual.

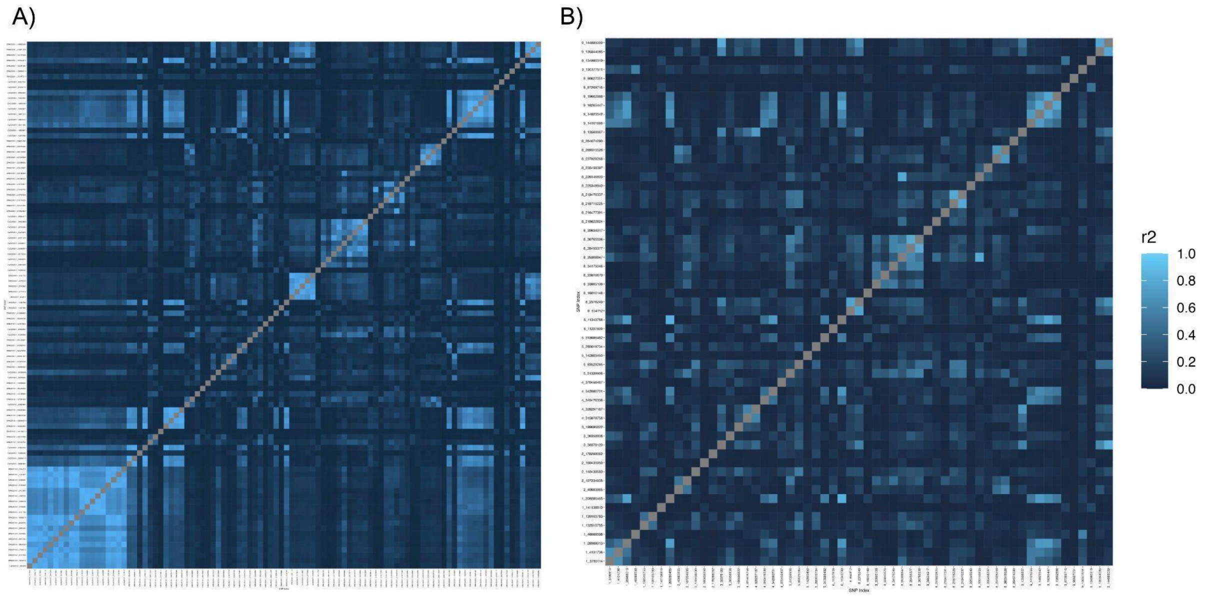


**Figure S6- Linkage disequilibrium measured using the correlation coefficient between SNPs strongly associated with precipitation in the GDM analysis.** Correlation between SNPs before (A) and after (B) the removal of highly correlated SNPs ($r^2 \geqq 0.8$) occurring on the same chromosome.

**Table S3- Model fit estimates for PRS models using unlinked SNPs.**

| | **Cubic** | | **Polynomial** | | **Linear** | |
|---|---|---|---|---|---|---|
| | $R^2$ | AIC | $R^2$ | AIC | $R^2$ | AIC |
| **Filtered (n= 59)** | 0.94 | -124.82 | 0.92 | -118.09 | 0.78 | -100.32 |

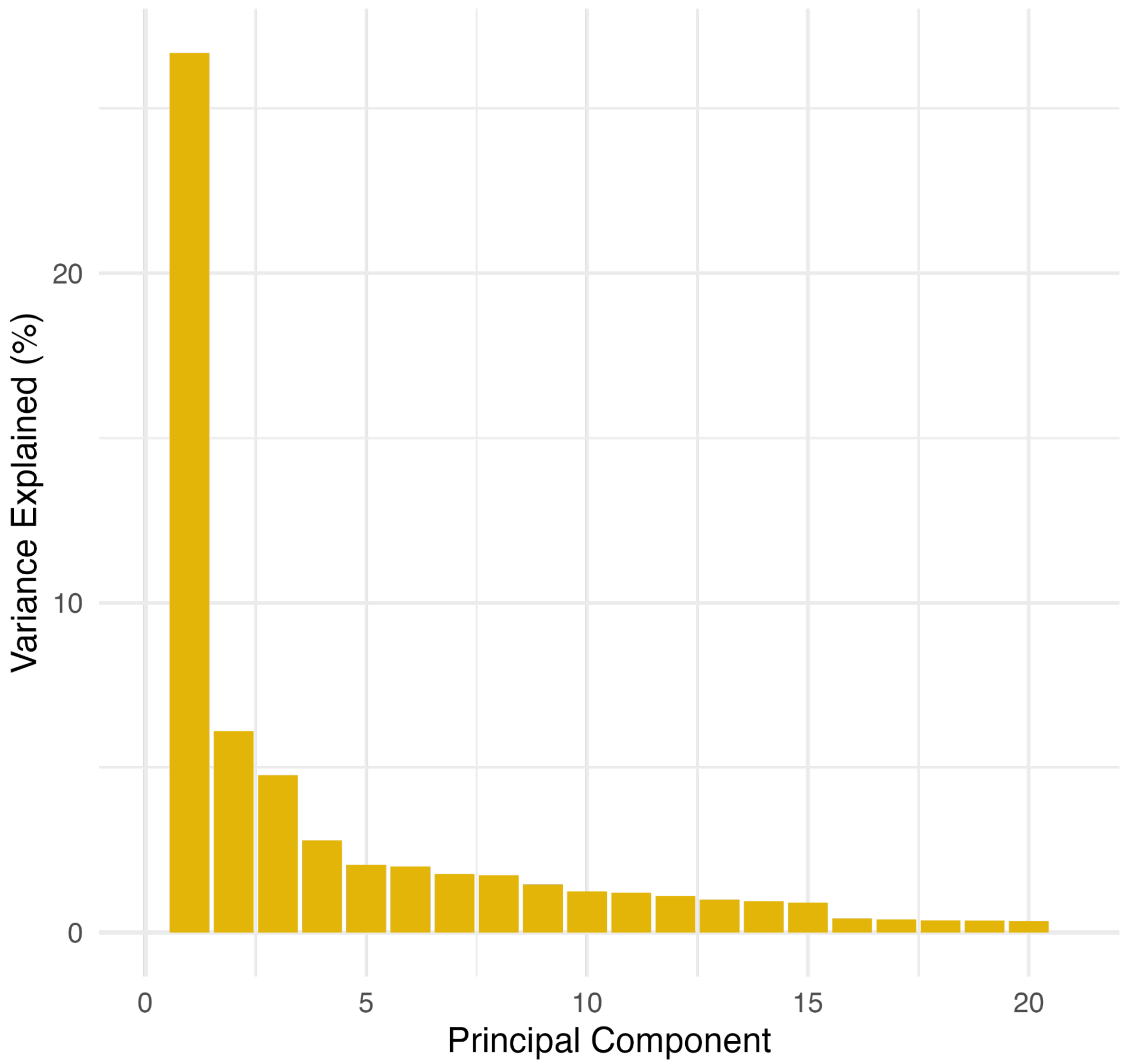


**Figure S7- Percentage variance explained by each PC of a PCA on individual genotype likelihoods.** PCA of individual genotype likelihoods (n=329) and the variance explained by each principal component.

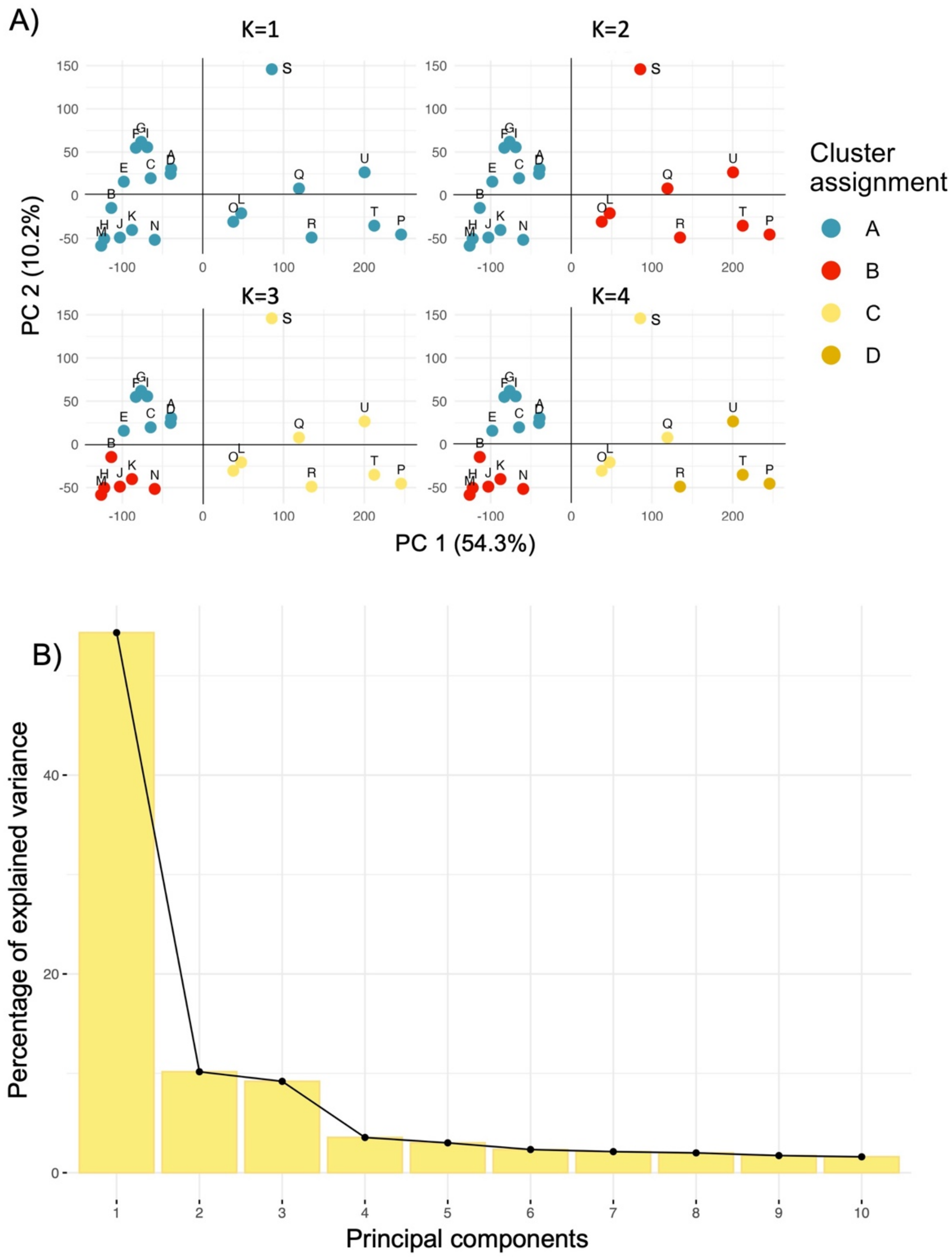


**Figure S8 - PCA and kmeans clustering of population allele frequencies.** (A) Clustering of 580,412 allele frequencies for each population for K 2 - 4 and (B) the variance explained by each principal component visualized.

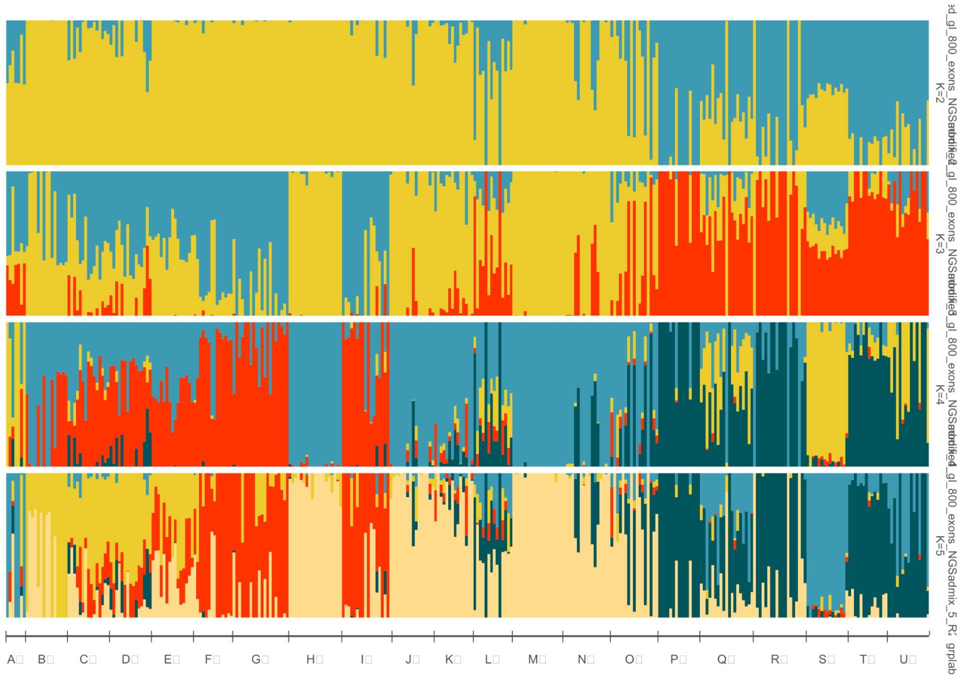


**Figure S9: STRUCTURE plot of individuals.** Letters A-U indicate populations, where A is the driest population in mean annual precipitation from 1970-2000 from WorldClim (Fick & Hijmans, 2017). Depicted are *K* 2-5, where each individual's (n = 329) genome is represented as a vertical bar partitioned into colored segments. These segments indicate the inferred admixture proportions, reflecting the estimated percentage of an individual's genome originating from each of the K identified genetic clusters.

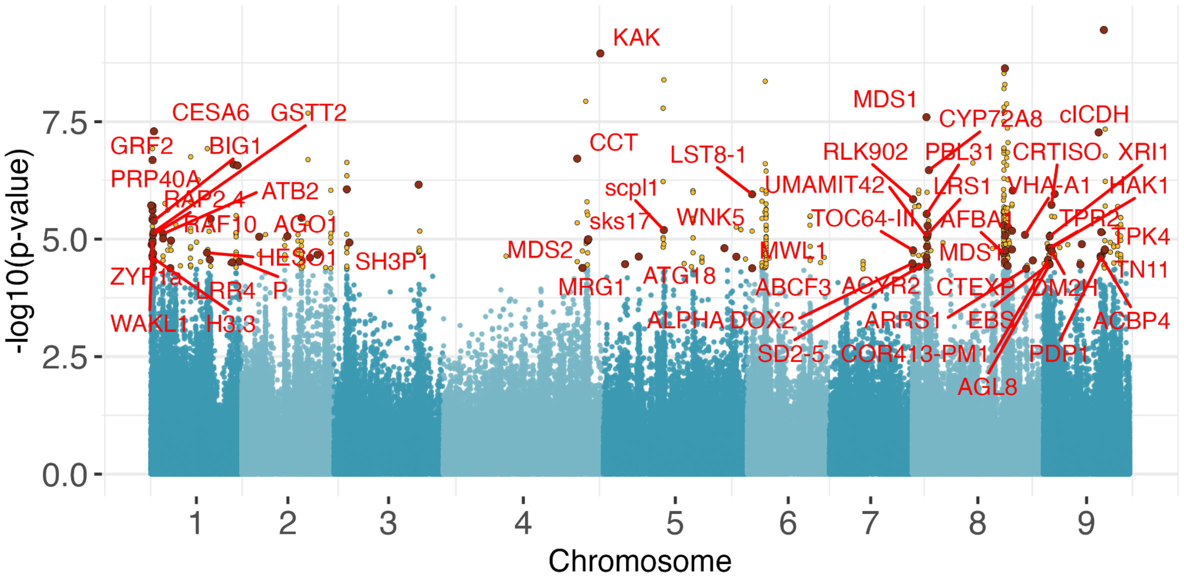


**Figure S10- Manhattan plot of SNPs identified as significantly associated with precipitation of the wettest month using LFMM.** All 580,412 SNPs are depicted. The 488 SNPs significantly associated with precipitation of the wettest month (yellow). SNPs with allelic turnover most strongly associated with precipitation and showing weak associations with temperature and geography are depicted as red points, and those with *Arabidopsis thaliana* orthologs are correspondingly labeled (n=55).

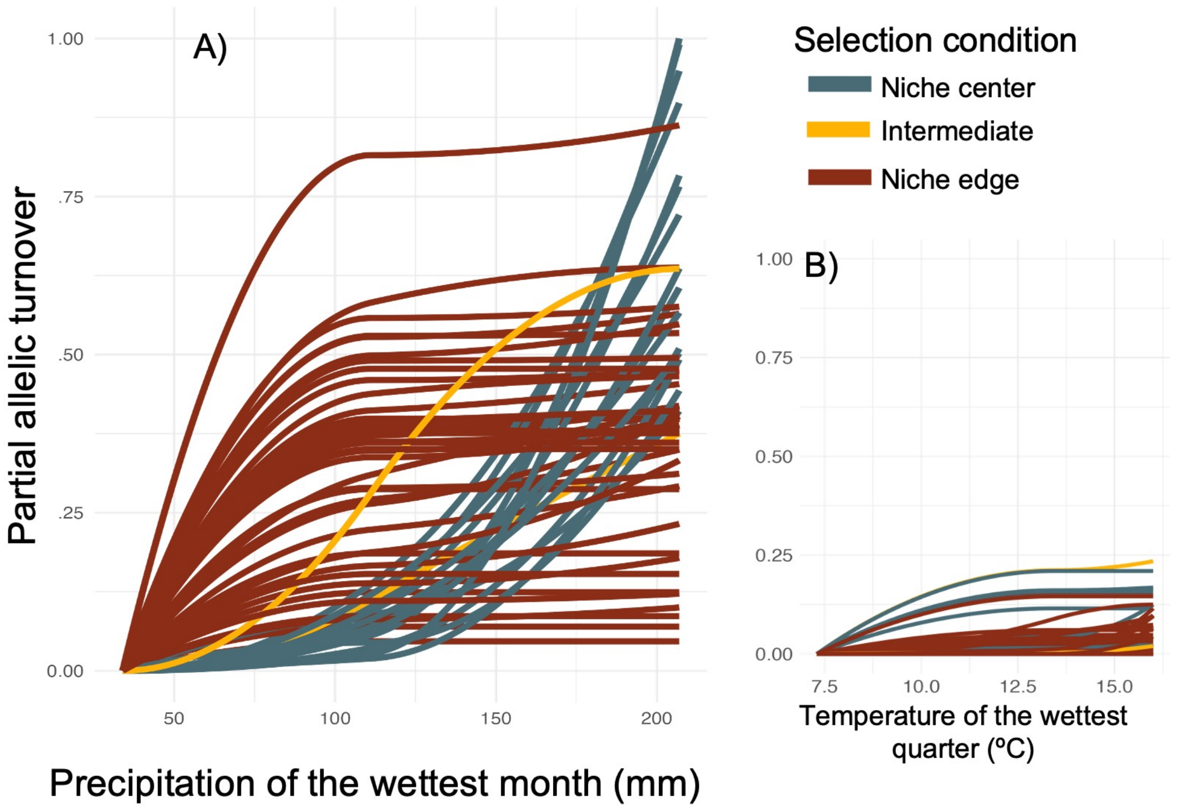


**Figure S11: Adaptive variation along the precipitation gradient for unlinked SNPs.** (A) Allelic turnover of unlinked candidate SNPs only (n=59). SNPs are classified according to their highest turnover at the arid niche edge (50-100 mm; red), intermediate (100-150 mm, yellow), and the transition to more mesic conditions (150-200 mm, blue). (B, C) Turnover for these same candidate SNPs visualized across geographic distance and temperature.

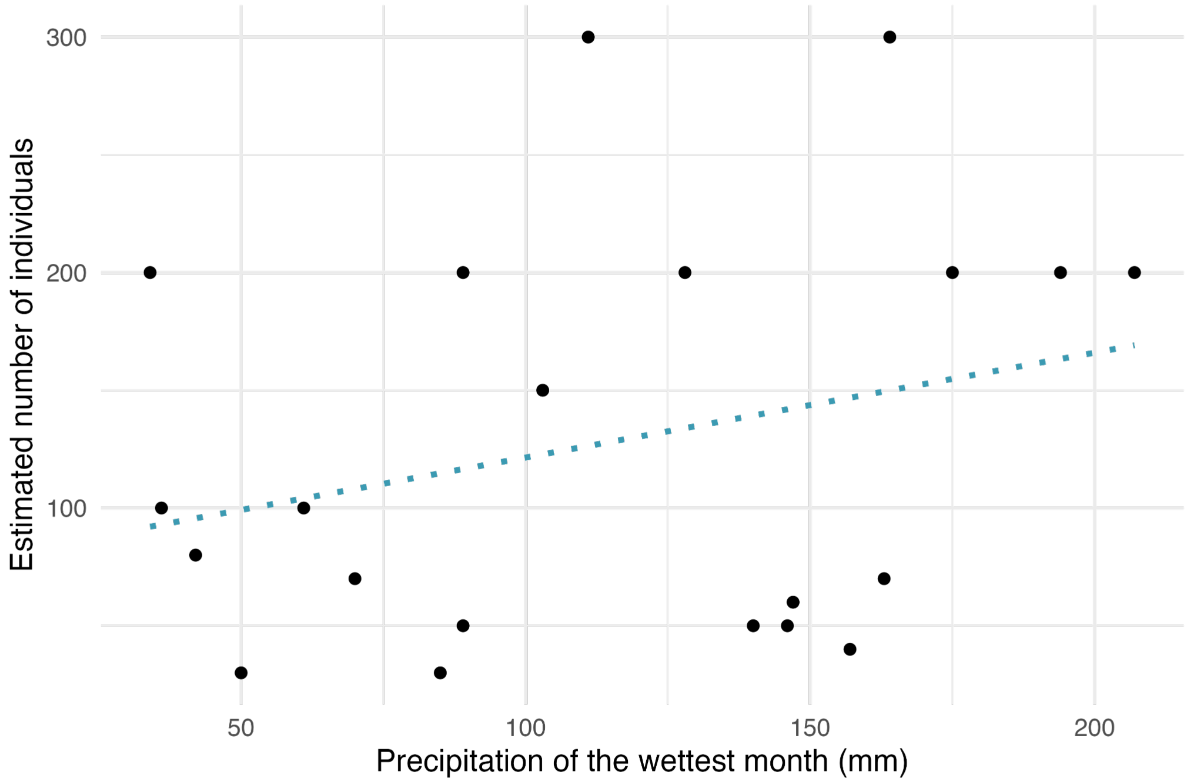


**Figure S12: Estimated population size for populations sampled in this study**. Approximate population size (number of adult plants) was visually estimated at the time of seed collection in the summer of 2020 for each population. A linear model (blue line) between precipitation of the wettest month and estimated number of individuals in the population is not significant ($P$ = 0.30; Adjusted $r^2$ = 0.026).

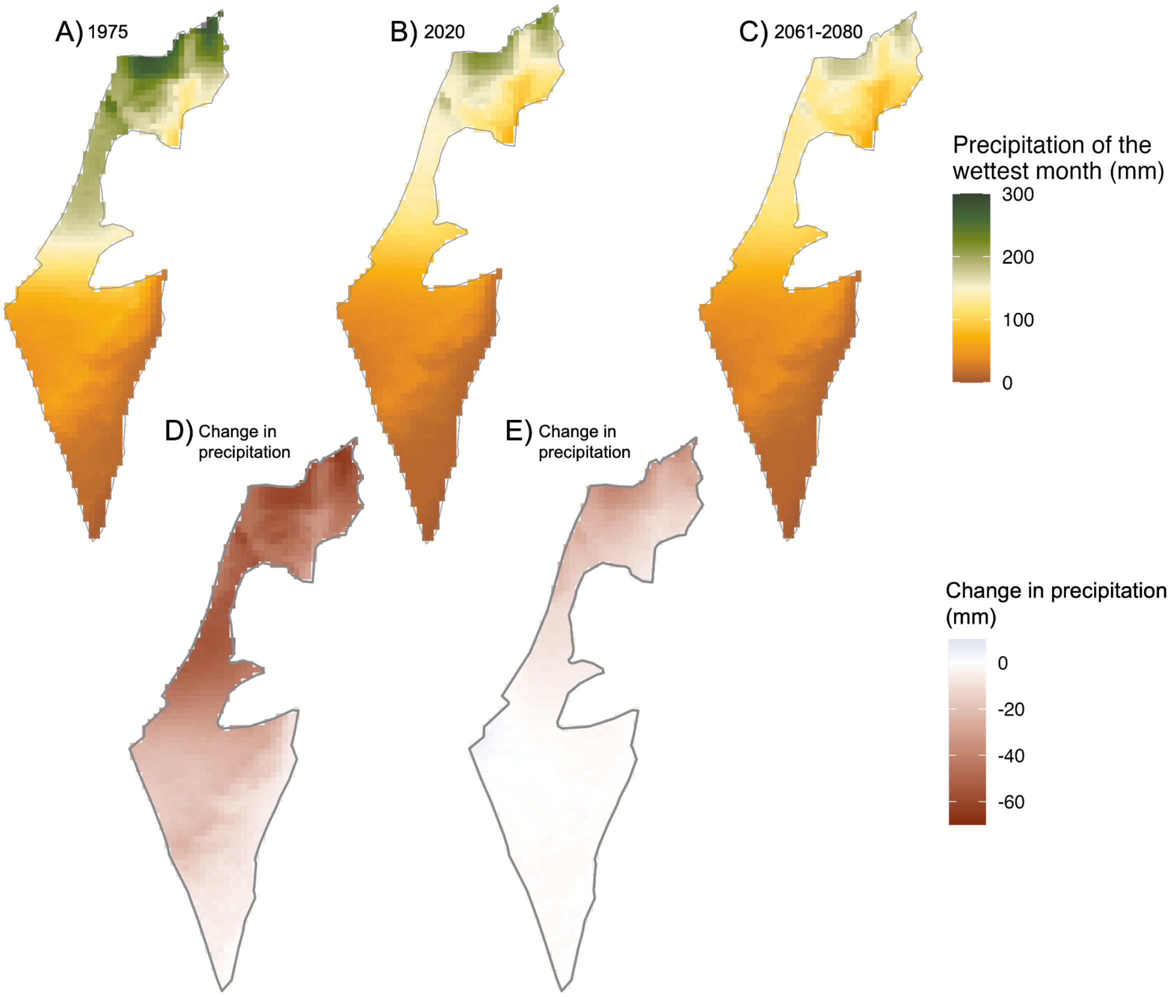


**Figure S13: Precipitation of the wettest month across Israel.** Precipitation values at a 2.5-arcminute resolution across Israel for (A) 1975, (B) 2020, and (C) the mean estimate for the period 2061-2080. (D) The change in precipitation of the wettest month between 1976 and 2020, and (E) the change in precipitation of the wettest month between 2020 and the mean predicted values for the period of 2061-2020.
